# PAF1C-driven restoration of RNAPII elongation after DNA damage occurs independently of transcription-associated histone mark deposition

**DOI:** 10.1101/2025.07.23.666359

**Authors:** Janne J.M. van Schie, Bram A.F.J. de Groot, Martijn S. Luijsterburg

## Abstract

DNA lesions block the progression of RNA polymerase II (RNAPII) during transcription, impeding gene expression and threatening genome integrity. When RNAPII stalls on transcription-blocking lesions, the transcription-coupled DNA repair pathway is activated to remove the DNA damage. Following DNA repair, efficient transcription restart depends on the PAF1 elongation complex (PAF1C). PAF1C contributes to deposition of transcription-associated histone marks, including H2B-K120_Ub_, H3K4me_3_ and H3K79me_2_. These marks are enriched at actively transcribed genes and have been associated with regulation of post-repair transcription restart. Here, we show that the H2B-K120 E3 ubiquitin ligase RNF20/RNF40, the H3K4-methyltransferase SET1/COMPASS complex, and the H3K79-methyltransferase DOT1L are dispensable for transcription restart. Moreover, levels of H2B-K120_Ub_ and H3K4me_3_ do not correlate with transcription restoration following DNA damage. Additionally, we observe that, unlike PAF1, the dissociable PAF1C subunit RTF1, while stimulating H2B-K120_Ub_ and H3K4me_3_, does not play a role in transcription restart. Together, these data suggest that transcription restoration after DNA damage is stimulated by the PAF1C elongation complex, independently of transcription-associated histone mark deposition.

## Introduction

The presence of DNA damage in the transcribed strand presents a major obstacle to transcription, leading to persistent stalling of RNA polymerase II (RNAPII) at bulky DNA lesions. This stalling is highly toxic and triggers a genome-wide transcriptional arrest. While DNA lesions lead to the direct stalling of elongating RNAPII *in cis*, there is also a regulated inhibition of transcription initiation *in trans* via the stress-induced repressor ATF3 and RNAPII degradation ^1-4^. Overcoming transcriptional arrest and restoring transcription after DNA repair is essential for maintaining proper gene expression and cellular homeostasis.

To remove transcription-blocking DNA lesions, cells rely on the transcription-coupled DNA repair (TCR) pathway. TCR is initiated by the coordinated recruitment of the TCR-specific proteins CSB, CSA, UVSSA and STK19 to DNA damage-stalled RNAPII assisted by elongation factor ELOF1 ^5-11^. Together, these factors promote the recruitment of the TFIIH complex to damage-stalled RNAPII, leading to either backtracking or displacement of RNAPII to facilitate access to downstream repair proteins. Subsequently, general NER proteins, such as XPA, RPA, and the endonucleases XPG and ERCC1-XPF, coordinate dual incision around the lesion to remove the damaged DNA strand, followed by gap fill DNA synthesis and ligation. Mutations in TCR components CSB and CSA cause Cockayne syndrome, a progeroid disorder characterized by severe developmental and neurological dysfunction ^12,13^.

While the removal of DNA lesions by TCR is essential, it is not sufficient for transcription restart. TCR involves displacement of RNAPII from the damaged DNA template ^14-17^, followed by transcription restart from gene promoters. Post-repair transcription restart requires the alleviation of ATF3-mediated transcriptional repression at gene promoters by CSB- and CSA-mediated degradation of ATF3 ^1,18^. Furthermore, chromatin remodeling has been implicated in post-repair transcriptional restart ^19^. While these observations suggest post-restart repair pathways are at least partially distinct from general transcription requirements, the precise mechanisms driving transcription recovery and their coordination with TCR-mediated repair remain to be fully elucidated.

The PAF1 subunit of the RNA polymerase II-associated factor complex (PAF1C) has emerged as a critical component in regulating transcriptional restart following genotoxic stress ^20^. The PAF1C, composed of subunits CTR9, PAF1, LEO1, CDC73, SKI8 and the loosely associated subunit RTF1, localizes to actively transcribed genes through its association with RNAPII to regulate transcription elongation, promoter-proximal pausing, and the deposition of histone modifications ^21-26^. Which functions of the PAF1C contribute to transcription restart and whether subunits other than PAF1 are required for restart is currently unknown.

PAF1C recruits the ubiquitin conjugase (E2) RAD6A (UBE2A) and the RING-type E3 ubiquitin ligase complex RNF20-RNF40 to RNAPII to co-transcriptionally mono-ubiquitylate histone H2B at lysine 120 (H2BK120_ub_) ^27,28^. In both yeast and human, this modification stimulates the subsequent deposition of H3K4 trimethylation (H3K4_me3_) by the SET1/COMPASS complex and H3K79 dimethylation (H3K79_me2_) by DOT1L ^28-33^. Thereby, PAF1C stimulates the deposition of histone marks H2BK120_Ub_, H3K4_me3_, and H3K79_me2_ ^34^ which are all associated with active transcription ^35,36^. H3K4_me3_ is mainly enriched close to promoters and correlates with transcription initiation and promoter proximal pause release ^37,38^. H2BK120_Ub_ and H3K79_me2_ are enriched in gene bodies of active genes, and have both been linked to transcription elongation and the DNA damage response ^39-41^. In addition, H2BK120_ub_, H3K4_me3_ and H3K79_me2_ have been associated with restoring post-repair transcription restart. UV-induced H3K4 methylation facilitates transcriptional reactivation in *C. elegans* ^42^, while DOT1L-mediated H3K79 methylation contributes to transcription restart in mouse cells ^43^. Furthermore, H2BK120_ub_ levels are reduced after UV damage ^20,44^. Together, these findings suggest a potential interplay between transcription-associated histone modifications and transcription recovery ^20^, but this has not been directly assessed in human cells.

In this study, we found that transcription-associated histone modifications H2BK120_Ub_, H3K4_me3_, and H3K79_me_ are dispensable for transcriptional recovery following genotoxic stress. Additionally, unlike PAF1, the dissociable PAF1C subunit RTF1, while stimulating H2B-K120_Ub_ and H3K4_me3_, does not play a role in transcription restart. Together, our work suggests that restoration of transcription following DNA damage depends on the ability of PAF1C to directly stimulate processive RNAPII elongation in a manner that is independent of the deposition of transcription-associated histone marks.

## Results

### DOT1L and H3K79 methylation are redundant for transcription restart following DNA damage

H3K79_me2_ is a histone mark enriched in gene bodies of actively transcribed genes and correlates with high RNAPII elongation rates ^41^. However, loss of H3K79_me2_ does not seem to impact transcription elongation rates ^45,46^, and only causes minor changes in gene expression ^47,48^. H3K79 methylation has been linked with transcription initiation ^46^, but the exact contribution of H3K79_me2_ in transcription remains unclear ^41^. Previous work has shown that H3K79_me2_ facilitates transcription restart after genotoxic stress in mouse cells, possibly by opening chromatin structure to allow reactivation of RNAPII by facilitating transcription initiation ^43^. Since differences in chromatin requirements for TCR between mouse and human cells have been observed before ^49^, we decided to test the dependence of transcription restart after DNA damage repair on H3K79_me2_. To do so, we inhibited DOT1L, the only known methyltransferase of H3K79, with DOT1L inhibitor Pinometostat (DOT1Li) in human RPE1-hTERT cells. Because of the lack of an obvious H3K79 demethylase ^41^, reduction of H3K79_me2_ seems instead to depend on histone replacement or dilution through cell division ^50^. Consequently, H3K79_me2_ has a relatively long half-life (multiple days) ^51^, which in itself makes it an unlikely candidate to dynamically respond to transcription-blocking DNA damage. To test the contribution of H3K79_me2_, we treated cells for 7 days with DOT1Li, which resulted in a complete loss of H3K79_me2_ as detected by western blot analysis (Fig 1A) and performed recovery of RNA synthesis (RRS) assays after UV-induced DNA damage. In all conditions, a drop of RNA synthesis was observed 3h after UV exposure. As expected, TCR-deficient CSB-KO cells were unable to recover RNA synthesis after 24h. In contrast, nascent RNA synthesis in wild-type (WT) cells treated with DOT1Li recovered to levels similar to untreated WT cells (Fig 1B). Because incomplete inhibition of H3K79 methylation or H3K79_me2_-independent DOT1L functions ^45^ cannot be excluded using DOT1Li, we also generated two clonal DOT1L-KO cell lines using two different crRNAs targeting exon 2 and exon 5, respectively. Isolated DOT1L-KO clones had no detectable DOT1L protein or H3K79_me2_ (Fig 1C), and showed comparable levels of nascent transcription in unirradiated cells (Fig 1D). Similar to the cells treated with DOT1Li, also DOT1L-KO cells recovered nascent transcription like WT cells 24h after UV irradiation (Fig 1E). These observations show that DOT1L and H3K79 methylation are dispensable for transcription restart following genotoxic stress in human RPE1 cells.

**Figure 1.**
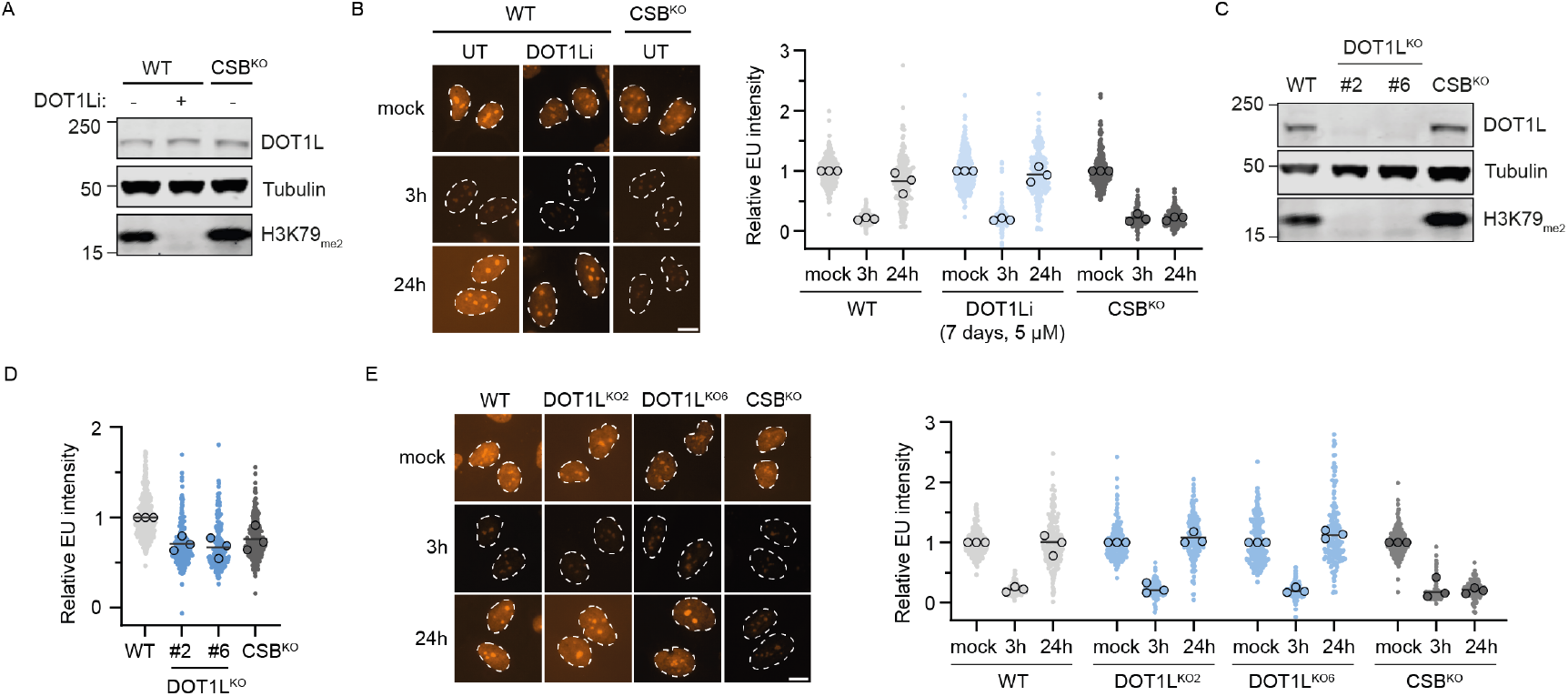
H3K79_me2_ is dispensable for transcription restart after DNA damage. **a**. Western blot with indicated antibodies of WT and CSB-KO cells untreated or treated with 5 μM DOT1Li (pinometostat) for 7 days. **b**. RNA recovery assay (RRS) of conditions from (a). Cells were EU labeling for 1h in unirradiated (mock) condition or 3 h and 24 h after 12 J/m^2^ UV. Cells depicted as individual data points. Circles are individual means of three biological replicates. Lines indicate means. Scale bar, 10 μm. **c**. Western blot of RPE1 cells with the indicated genotype. **d**. Nascent RNA synthesis in nonirradiated cells relative to WT cells of cell from (c). **e**. RRS as in (b) of cells from (c). Scale bar, 10 μm.

### The SET1 complex and H3K4 trimethylation are dispensable for transcription restart

H3K4_me3_ is enriched near promoters and is believed to aid transcription initiation and release of promoter proximally paused RNAPII ^37,38^. Consequently, RNAPII restart from promoters after DNA damage is repaired may therefore depend on H3K4_me3_. The deposition of H3K4_me3_ is mediated by the SET1 histone methyltransferase complexes. The H3K4 methyltransferase activities of all SET1 complexes depend on the shared WRAD complex, consisting of core subunits, WDR5, RBBP5, ASH2L and DPY30 ^52^. To test H3K4_me3_ involvement in transcription restart after UV, we depleted the shared core subunit RBBP5. Because the SET1 complex is essential for cell viability, likely due to its central role in transcriptional regulation ^52^, we were unable to generate viable RBBP5-KO clones. Instead, we acutely depleted RBBP5 using CRISPR/Cas9 and performed our assays after 5 days, when cells are still viable but have completely lost detectable levels of H3K4_me3_ (Fig 2A). We subsequently measured nascent transcription levels by 5-ethynyl-uridine (5-EU) incorporation while at the same time co-staining for H3K4_me3_ (Fig 2B). This approach confirmed the loss of H3K4_me3_ (defined as H3K4_me3_ levels lower than 0.2 relative to WT, cells depicted as pink dots) in around 90% of crRBBP5-transfected cells. However, near-complete loss of H3K4_me3_ in RBBP5-depleted cells did not affect the levels of nascent transcription when compared to control cells (Fig 2C). We next performed RRS assays while also co-staining for H3K4_me3_ (Fig 2D-E). While cells acutely depleted for TCR factor CSA were unable to recover RNA synthesis 24h after UV irradiation, RBBP5-depleted cells recovered like WT cells (Fig 2D). The levels of H3K4_me3_ were not decreased at 3h after UV irradiation in cells transfected with negative control (NC) or CSA crRNAs when nascent transcription is reduced by ∼80% (Fig 2D). Also in CSA-depleted cells at 24h, which fail to recover transcription and remain to have very low nascent RNA synthesis levels, this does not result in differences in H3K4_me3_ levels. Finally, RBBP5-depleted cells at 24h after UV, while being devoid of H3K4_me3,_ showed normal recovery of nascent transcription (Fig 2D). Together, this shows that the SET1 complex and H3K4_me3_ are not require for the immediate restart of transcription following the repair of transcription-blocking DNA damage.

**Figure 2.**
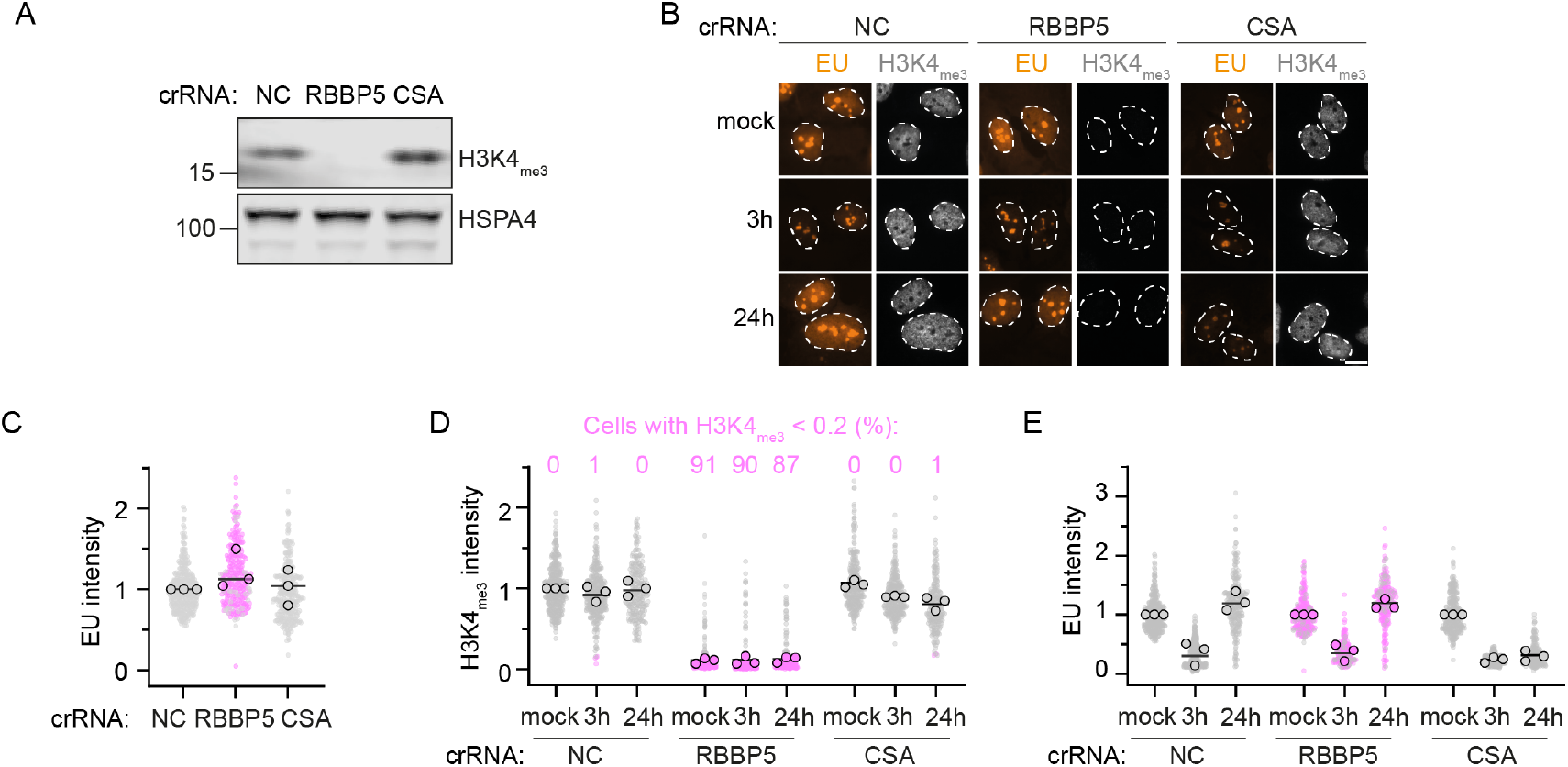
H3K4_me3_ is dispensable for transcription restart after DNA damage. **a**. Western blot of RPE1 cells 5 days after Cas9 induction and transfection with indicated crRNAs. **b**. Representative images of cells labelled for 1h with EU, unirradiated, 3h and 24h after 12 J/m^2^ UV, 6 days after transfection with indicated crRNAs. Scale bar, 10 μm. **c**. Nascent EU incorporation in nonirradiated cells relative to WT cells of cells transfected with indicated crRNAs. **d**. Quantification of H3K4_me3_ levels in conditions from (b), normalized to crNC mock condition. Individual data points represent cells, circles represent means from biological replicates. Pink dots represent cells with relative H3K4_me3_ levels below 0.2, numbers indicate percentage of cells with relative H3K4_me3_ levels below 0.2 **e**. Quantification of EU levels in conditions from (b), normalized to the mock condition per crRNA.

### H2BK120 ubiquitination and RNF20 are not essential for transcription restart

H2BK120_Ub_ is a co-transcriptionally deposited histone mark that is enriched within gene bodies of actively transcribed genes ^28^. H2BK120_Ub_ has been ascribed multiple functions in the DNA damage response ^40^. Furthermore, levels of H2BK120_Ub_ were shown to decrease following UV-induced DNA damage ^20,44^, suggesting a potential correlation, although not necessarily causal, between transcription restart after UV and this histone mark. H2BK120_Ub_ is deposited by the E2 conjugase RAD6 together with the heterodimeric E3 ligase RNF20-RNF40, which is essential for cellular viability. To test whether H2BK120_Ub_ contributes to recovery of transcription after genotoxic stress, we acutely depleted RNF20 using crRNF20 transfection together with Cas9 activation, which resulted in a strong reduction in H2B-K120_Ub_ within 5 days following transfection(Fig 3A-C). More than 70% of RNF20-depleted cells had H2BK120_ub_ levels of less than 0.2 relative to cells transfected with a control (NC) crRNA (cells depicted as purple dots). Cells depleted for RNF20 were still able to normally recover RNA synthesis after UV (Fig 3D), suggesting that H2BK120_ub_ is dispensable for transcription recovery after UV. In line with previous results ^20,44^, H2BK120_Ub_ levels dropped after UV irradiation, dropping concomitantly with the reduction in transcription observed at the 3h post-UV timepoint (Fig 3C). However, when transcription recovered in control cells 24h after UV irradiation, H2BK120_Ub_ levels were still not fully recovered.

**Figure 3.**
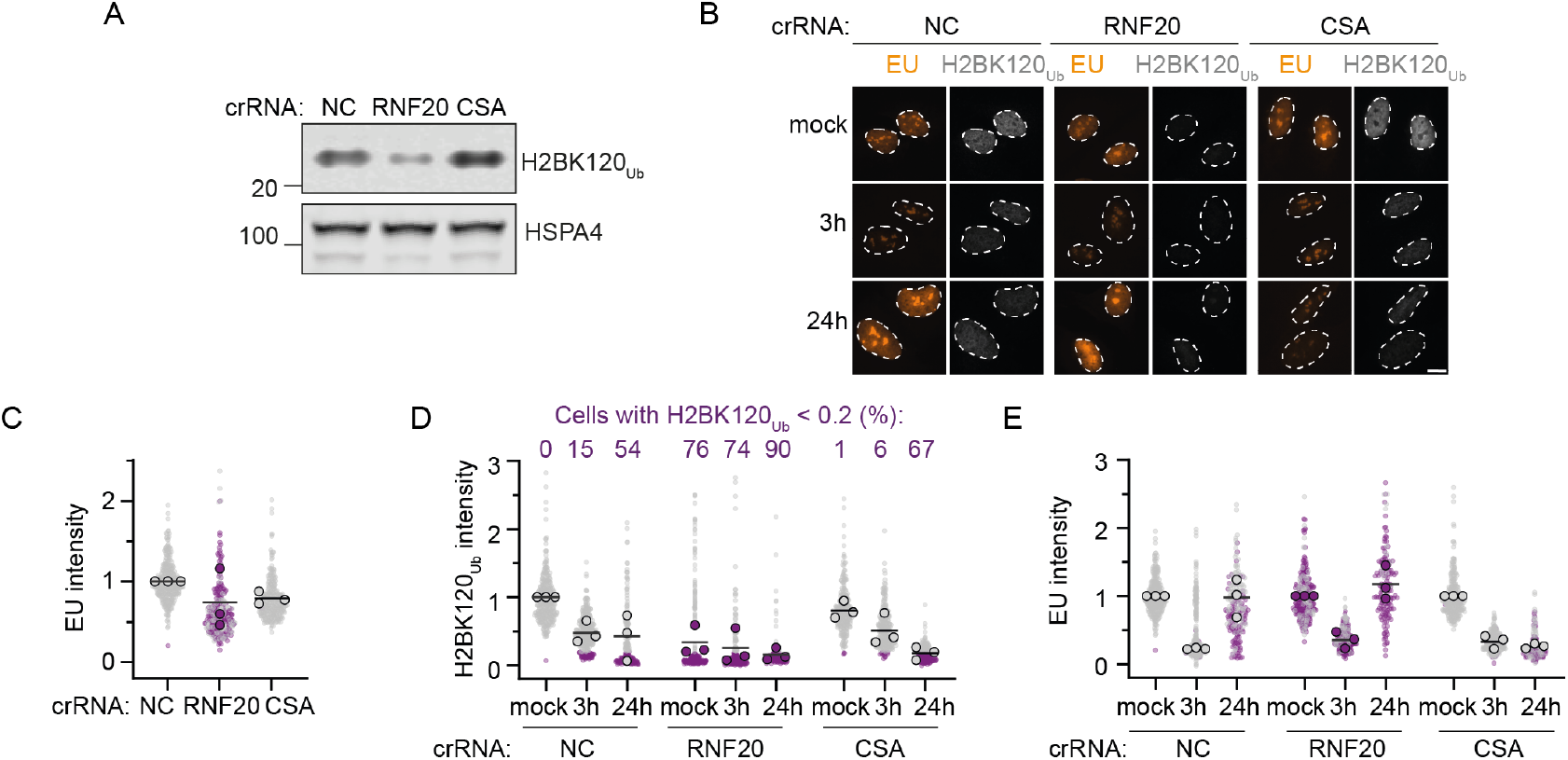
H2BK120_Ub_ is dispensable for transcription restart after DNA damage. **a**. Western blot of RPE1 cells 5 days after Cas9 induction and transfection with indicated crRNAs. **b**. Representative images of cells labelled for 1h with EU, unirradiated or 3 h and 24h after 12 J/m^2^ UV, 6 days after transfection with indicated crRNAs. Scale bar, 10 μm. **c**. Nascent EU incorporation in nonirradiated cells relative to WT cells of cells transfected with indicated crRNAs. **d**. Quantification of H2B-K120_Ub_ levels in conditions from (b), normalized to crNC mock condition. Individual data points represent cells, circles represent means from biological replicates. Pink cells are cells with relative H2B-K120_Ub_ levels below 0.2, numbers indicate percentage of cells with relative H2B-K120_Ub_ levels below 0.2. **e**. Quantification of EU levels in conditions from (b), as in (c) but normalized to mock condition per crRNA.

Together, this shows that H2BK120_Ub_ and RNF20 are not essential to restart transcription after UV-induced DNA damage. Rather, these findings suggest that H2BK120_Ub_ levels drop as a result of decreased transcription after UV, while restoring H2BK120_Ub_ levels after UV lags behind RNAPII elongation and is substantially slower than the actual restart of transcription.

### Core PAF1C subunits PAF1 and CTR9, but not subunit RTF1, is required for transcription restart

Since both PAF1 and RTF1 were found to physically interact with RAD6A/RNF20-RNF40 to stimulate H2BK120 ubiquitylation ^27,28,33^ and in turn H2BK120_ub_ stimulates H3K4_me3_ and H3K79_me2_ deposition, we decided to test levels of these marks after depletion of PAF1 and RTF1. Depletion of PAF1 and more prominently for RTF1 resulted in a pronounced drop in H2B-K120_ub_ levels (Fig. 4A-B). In addition, levels of H3K4_me3_ were also strongly decreased, particularly in RTF1-depleted cells. RTF1 or PAF1 depletion had a less pronounced effect on the H2BK120_ub_-stimulated histone mark H3K79_me2_ ^28-33^ (Fig. 4A-B), possibly due to the long half-life of this mark ^53^ or H2B-K120_ub_-independent H3K79 methylation pathways.

**Fig 4.**
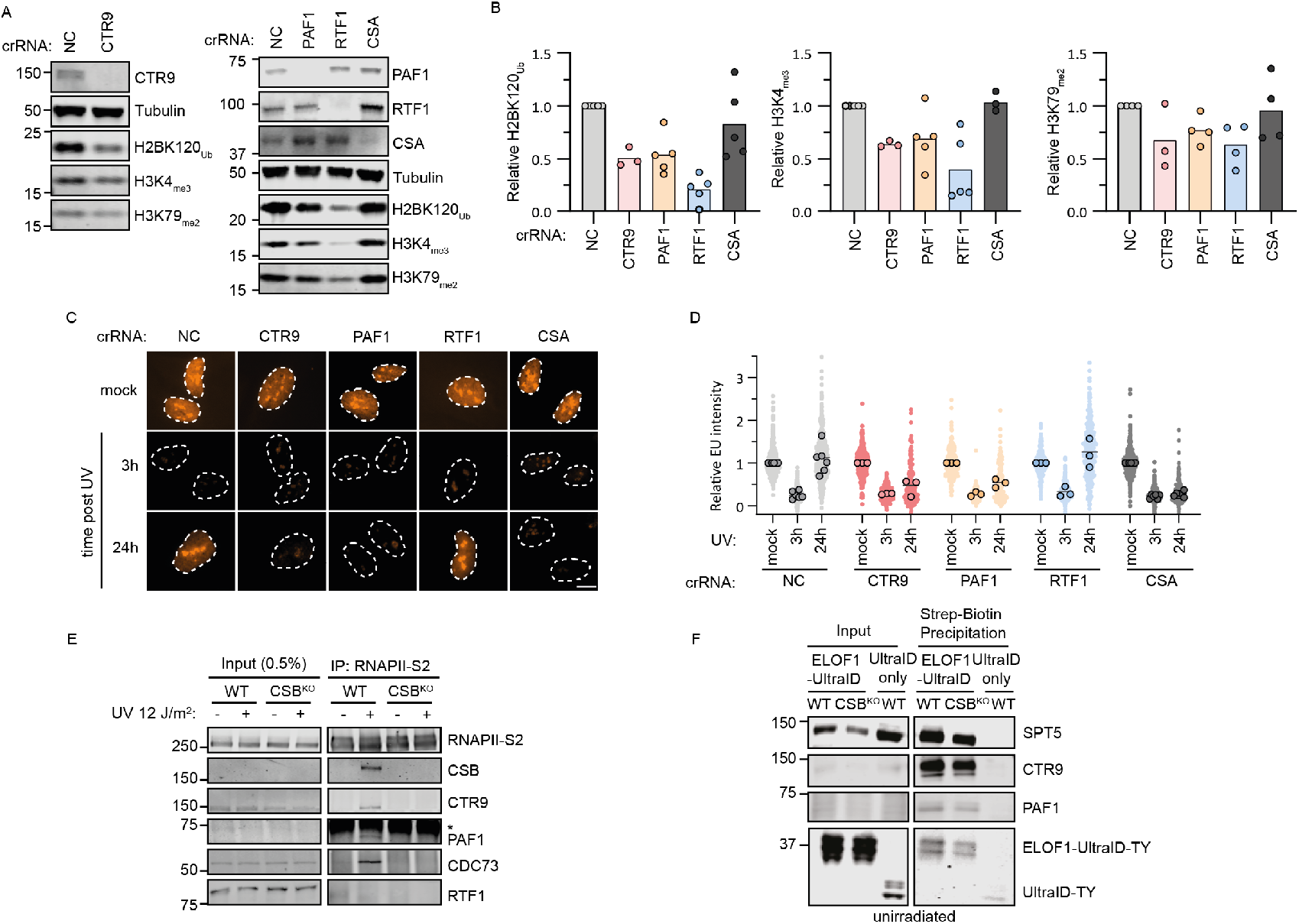
RTF1 is not required for transcription restart. **a**. Western blot of RPE1 cells 5 days after Cas9 induction and transfection with indicated crRNAs. **b**. Quantifications of western blots of H2B-K120Ub (left), H3K4me3 (middle) and H3K79me2 (right) levels, normalized to loading control tubulin and relative to crNC. **c**. Representative images from RRS assay of cells 6 days after transfection with indicated crRNAs. **d**. RRS of conditions from (c). Cells were EU labelled for 1h in cells unirradiated, 3 h and 24 h after 12 J/m^2^ UV. Cells are depicted as individual data points. Circles are individual means of three biological replicates. Lines indicate means. Scale bar, 10 μm. **e**. RNAPII-S2 co-immunoprecipitation in unirradiated (-) or 1h after 12J/m^2^ UV irradiation in WT and CSB^KO^ cells. **f**. ELOF1-UltraID proximity biotin-labelled proteins were streptavidin precipitated followed by detection with western blot. ELOF1-T2A-UltraID (right lane) is a negative control.

Previous work revealed that the PAF1C subunit PAF1 is required for transcription restart after UV irradiation ^20^. Furthermore, it was found that RNAPII stably interacts with five PAF1C subunits (CTR9, PAF1, CDC73, PAF1 and SKI8), but not with RTF1, in response to UV-induced DNA damage ^20^. While PAF1 and RTF1 are part of the same complex in yeast, RTF1 does not stably associate with PAF1C in human cells ^22,34,54^. Therefore, we hypothesized that these subunits may have different contributions to transcription restart. In line with previous results using siRNAs or a degron system to deplete PAF1 in U2OS cells ^20^, we found that acute knockout of PAF1 by transfection of crPAF1 in RPE1 cells impaired transcription restart after UV irradiation (Fig 4C-D). To test whether this is a general feature of the PAF1 core complex, we performed an acute knockout of another core subunit, CTR9, which similarly impaired transcriptional restoration (Fig. 4C–D). In contrast, acute knockout of RTF1, which was confirmed by western blot analysis, did not affect transcription recovery (Fig 4C-D), suggesting that RTF1 is dispensable for this process. Together, our findings show that core PAF1C proteins, but the not dissociable subunit RTF1 drives transcription restoration after UV in a manner that is not mediated via H3K79_me2_, H2B-K120_ub_ or H3K4_me3_ deposition.

To study the relationship between PAF1C binding to RNAPII in more detail, we employed co-immunoprecipitation, which is better suited for detecting DNA damage-specific interactions, and proximity labeling, which is more appropriate for capturing transient binding of elongation factors to RNAPII during unperturbed transcription. Co-immunoprecipitation of RNAPII-S2 from UV-irradiated cells showed that the interaction of PAF1, but not RTF1, with RNAPII is enhanced by UV damage in a CSB-dependent manner (Fig 4E). Our co-immunoprecipitation approach fails to detect PAF1 or RTF1 interactions with RNAPII in undamaged conditions, while these are well described interactions ^34^. As mentioned, this is likely because the interactions are too transient and therefore not detected using our IP approach. Proximity labelling methods, such as those using biotin ligase UltraID, allow detection of transient protein-protein interactions ^55^. To study transient interactions of elongating RNAPII, we endogenously fused UltraID to elongation factor ELOF1 using a knock-in approach ^5,7^. As a control, we knocked UltraID into the ELOF1 locus, flanked by a T2A peptide that causes ribosome skipping during translation and thereby produces two separate proteins, ELOF1 and UltraID, from a single mRNA transcript. Indeed, this proximity-labelling approach showed that PAF1C interacts with elongating RNAPII in the absence of DNA damage (Fig 4F), as expected, while no interactions were detected with the UltraID only control. Importantly, the interaction of PAF1C with elongating RNAPII is independent of CSB (Fig 4F). Together, these findings indicate that DNA damage induces a CSB-dependent stabilization in the interaction between PAF1C and RNAPII, which is likely central to the transcription restart function of PAF1C.

Together, we demonstrated that the PAF1C elongation complex contributes to transcription restart after DNA damage, independently of the RTF1 subunit and in a manner that does not correlate with or depend on transcription-associated histone marks H3K79_me2_, H2B-K120_ub_ and H3K4_me3_.

## Discussion

Our study demonstrates that initial post-repair transcription restart is independent of histone modifications H2B-K120_Ub_, H3K4_me3_ and H3K79_me2_. While these marks correlate with active transcription, they are not strictly required for transcription ^45 41 56 57 58^. Our data indicates that RNA synthesis can happen in the absence of these marks, both in unchallenged and post-repair conditions.

Previous studies in model systems other than human cells have shown that transcription restart after DNA damage partially depends on methylation of either H3K4 and H3K79. In *C. elegans*, H3K4 methylation and the SET1/COMPASS complex facilitate transcription restart after DNA damage^42^, whereas our findings show that neither H3K4_me3_ nor the SET1/COMPASS complex is required for transcription recovery in human cells. In line with our findings in human cells, in *C. elegans* only H3K4_me2,_ but not H3K4_me3_ dynamically changes in response to DNA damage ^42^. H3K4_me2_ and H3K4_me3_ have different dynamics, genomic distributions and have at least partially non-overlapping functions ^38,57,59^. Furthermore, H3K4 methyltransferases other than the SET1/COMPASS complex can contribute to H3K4_me2_ deposition ^42^, potentially explaining interspecies differences. Moreover, in the *C. elegans* experimental set-up, L1 larvae are irradiated and followed during development until adulthood, suggesting that H3K4_me2_ deposition may be a specific developmental response that we do not capture in cultured human cells ^42^.

In mouse embryonic fibroblasts, DOT1L-mediated H3K79 methylation has been reported to contribute to transcription restart ^43^, whereas we find no requirement for this modification in human cells. Similar to previously observed differences in TCR requirements between mouse and human cells ^49^, these findings may reflect interspecies differences in chromatin prerequisites for transcription recovery. We also note that DOT1L-deficient mouse cells were generated using a gene-trap insertion approach, and that the transcription restoration phenotype in these cells was not rescued by re-expression of DOT1L ^43,60^. This would be a critical control to rule out off-target effects caused by the gene-trap insertion. A previous report in human cells identified a role for DOT1L in transcriptional restart at sites of transcription-replication conflicts ^61^, suggesting dependency on DOT1L in transcription restart is damage and location specific.

Post-repair transcription restart can occur without histone mark deposition. Both PAF1 and CTR9, core subunits of the PAF1C transcription elongation complex are essential for efficient transcription restart, while the dissociable subunit RTF1 is dispensable. This finding is in line with the observation that RTF1 dissociates from RNAPII before CSB binding *in vitro* ^62^, while the other five subunits are stabilized on RNAPII after DNA damage *in vivo* ^20^. Moreover, considering that RTF1 is essential for PAF1C-dependent histone modifications, this already suggests that the role of PAF1C in transcription recovery is independent of the deposition of these marks. Interestingly, RTF1 does contribute to RNAPII velocity ^22^, suggesting that not all transcription elongation factors are required for transcription restart. PAF1 may influence transcription restart via regulation of (RTF1-independent) transcription elongation ^21,22,63^ or promoter-proximal pause release ^23,24,26^. In addition, PAF1C physically interacts with CSB ^20,64^, and the RNAPII interaction of PAF1 depends on CSB, suggesting this is a dedicated, TCR-specific pathway to activate post-repair transcription.

In conclusion, our study provides evidence that transcription recovery following genotoxic stress depends on PAF1C, but is independent of transcription-associated histone modifications and RTF1. It would be interesting to further dissect differences in specific requirements of post-repair restart compared to those of general transcription.

## Acknowledgments

MSL laboratory was supported by the European Research Council Consolidator Grant STOP-FIX-GO (grant agreement No 101043815) and the Netherlands Scientific Organization (NWO) Vici grant (VI.C.212.005). The funders had no role in study design, data collection and analysis, the decision to publish, or the preparation of the manuscript.

## Data Availability

This published article (and its supplementary information files) includes all data generated or analyzed during this study.

## Competing interests

The authors declare no competing interests.

## Author contribution

JJMvS generated knockouts; performed RRS experiments, immunofluorescence for histone marks, western blots for knockouts and histone marks, and co-immunoprecipitation; prepared figures; and wrote the paper. BAFJdG generated and validated ELOF1-UltraID cells and controls, performed proximity labeling coupled with western blotting, and prepared figures. MSL supervised the project, acquired funding, and wrote the paper.

## Methods

### Cell lines and cultures

The RPE1 cell lines were cultured at 37°C in an atmosphere of 5% CO_2_ in DMEM GlutaMAX (Thermo Fisher Scientific) supplemented with penicillin/streptomycin (Sigma), and 8% fetal bovine serum (FBS; Bodinco BV or Thermo Fischer Scientific (Gibco)).

### Generation of knockout cells

Parental RPE1-hTERT cells stably expressing inducible Cas9 (iCas9) that are also knockout for TP53 and the puromycin-N-acetyltransferase PAC1 gene were described previously (referred to as RPE1-iCas9). To generate knockouts in RPE1-iCas9, one day after induction of Cas9 by addition of 200 ng/ul doxycline (Clontech, 8634-1), cells were transfected with 10nM crRNA:tracrRNA duplexes using 1:1000 Lipofectamine RNAimax transfection reagent (Invitrogen, 13778150). crRNAs in Supplementary Table 2. Two days after transfection, single cells were plated by limiting dilution. After single cell clone expansion, clones were selected by western blot and Sanger sequencing using the oligos listed in Supplementary Table 4.

### Western blotting

Total cell lysates were harvested by scraping cells in the Laemmli-SDS sample buffer. Chromatin fractions were obtained by lysing cell pellets on ice in EBC-1 buffer (50 mM Tris [pH 7.5], 150 mM NaCl, 2 mM MgCl2, 0.5% NP-40, and protease inhibitor cocktail (Roche)) for 20 min at 4^°^ C, on a rotating wheel followed by centrifugation and removal of the supernatant. Chromatin pellets were resuspended in the Laemmli-SDS sample buffer. Total cell lysates or chromatin fraction samples were boiled for 10 min at 95°C. Proteins were separated on Criterion™ XT Tris-Acetate 3–8% Protein Gels (BioRad, #3450131) in Tris/Tricine/SDS Running Buffer (BioRad, #1610744) or on Criterion Xt bis-tris 4-12% gels in MOPS running buffer. Then, blotted onto PVDF membranes (IPFL00010, EMD Millipore) in Tris/glycine blotting buffer (0.025 M Tris, 0.192 M glycine) with 20% methanol. Membranes were blocked with 5% fat-free milk in PBS with 0.1 % Tween-20 for 1 h at room temperature. Membranes were then probed with indicated antibodies in 5% fat-free milk in PBS with 0.1 % Tween-20 (Antibodies are listed in Supplementary Table 5). Proteins were stained with fluorochrome-conjugated secondary antibodies and were detected on an Odyssey CLx system and Image Studio software (Li-Cor).

### Recovery of RNA synthesis (RRS)

After irradiation with UV-C light (12 J/m2), cells were allowed to recover for indicated times followed by pulse labelling with 400 μM 5-ethynyl-uridine (EU; Jena Bioscience) for 1 h. Next, cells were medium chased with DMEM without supplements followed by fixation in 3.7% paraformaldehyde in PBS for 15 min. Cells were permeabilized with 0.5% Triton X-100 for 10 min and blocked in 1.5% bovine serum albumin (BSA, Thermo Fisher) in PBS. Nascent EU labelled RNA was visualized by click-it chemistry, by incubating the cells with 60 μM Atto azide-Alexa 594 (Atto Tec), 4 mM copper sulfate (Sigma), 10 mM ascorbic acid (Sigma) and 0.1 μg/mL DAPI in 50 mM Tris-buffer pH 8 for 1 h. After washing extensively with PBS, slides were mounted in Polymount (Brunschwig).

### Microscopic analysis of fixed cells

Images of fixed samples were acquired on a Zeiss AxioImager M2 widefield fluorescence microscope equipped with 63x PLAN APO (1.4 NA) oil-immersion objectives (Zeiss) and an HXP 120 metal-halide lamp used for excitation. Fluorescent probes were detected using the following filters for DAPI (excitation filter: 350/50 nm, dichroic mirror: 400 nm, emission filter: 460/50 nm), Alexa 555/594 (excitation filter: 545/25 nm, dichroic mirror: 565 nm, emission filter: 605/70 nm), or Alexa 647 (excitation filter: 640/30 nm, dichroic mirror: 660 nm, emission filter: 690/50 nm). Images were recorded using ZEN 2012 software (blue edition, version 1.1.0.0) and analyzed in Image J (1.48v).

### Generation of ELOF1-UltraID cells

To generate endogenous knock-ins of UltraID-TY or T2A-UltraID-TY into the ELOF1 locus, RPE1-iCas9 WT and CSB-KO (1-15) cells were cultured in 6 cm plates prior to transfection. To promote homology-directed repair, cells were pretreated with PolQ inhibitor (ART558, MCE, 10 μM) and DNA-PK inhibitor (NU7441, MCE, 2 μM) 24 hours before transfection. RPE1 WT or CSB-KO (1-15) cells were then transfected with plasmids encoding a single guide RNA (sgRNA) targeting ELOF1, a Cas9-GFP construct (pML#187), and knock-in plasmids containing 1 kb homology arms targeting the ELOF1 locus, either ELOF1-UltraID-TY-T2A-Puromycin (pML#355) or T2A-UltraID-TY-T2A-Puromycin (pML#385), using 1 μg/μl PEI (Merck) in OptiMEM (Life Technologies, 51985034), followed by overnight incubation. Both plasmids (pML#355 and pML#385) contain a puromycin resistance cassette flanked by ∼1 kb homology arms to enable selection. After overnight transfection, cells were cultured for 2–3 days in standard DMEM supplemented with PolQ (10 μM) and DNA-PK (2 μM) inhibitors. GFP-positive cells were subsequently sorted by FACS and seeded. Single clones were isolated, transferred to individual culture plates, and selected for ELOF1-UltraID knock-ins using 1 μg/ml puromycin. Genomic DNA was extracted, and successful knock-in of UltraID or T2A-UltraID at the ELOF1 locus was confirmed by Sanger sequencing following nested PCR amplification of genomic DNA.

### Proximity labeling and streptavidin-based enrichment of biotinylated proteins

Cells were seeded and grown in 15 cm dishes until reaching 80% confluency. They were then incubated with 50 μM biotin for 15 minutes, followed by a 15-minute starvation in plain DMEM medium (Gibco, 31966-047). Subsequently, cells were collected by trypsinization and resuspended in 1× PBS at 4 °C. Immediately after resuspension, cells were pelleted by centrifugation at 1500 rpm and washed with 1× PBS. Cell pellets were snap-frozen in liquid nitrogen and stored at –80 °C until pull-down. Cells were lysed at 4 °C for 20 minutes using 1 ml of lysis buffer (50 mM Tris [pH 7.5], 150 mM NaCl, 0.1% deoxycholate, 2 mM MgCl_2_, 0.5% NP-40, and protease inhibitor cocktail [Roche]) supplemented with 500 U/ml benzonase (Novagen). After lysis, the volume was adjusted to 2.5 ml with lysis buffer. To remove free biotin and cell debris, the protein lysate was processed using a G25/PD-10 size exclusion column (Cytiva, 17085101) according to the manufacturer’s protocol. The resulting 3.5 ml protein eluate was incubated overnight at 4 °C with 30 μl streptavidin–sepharose beads (Merck Millipore, 69203-3) and 1000 U/ml benzonase to capture all biotin-labeled proteins. After six washes with washing buffer (50 mM Tris [pH 7.5], 300 mM NaCl, 0.1% deoxycholate, 1 mM EDTA, 0.5% NP-40), samples were boiled in 2× Laemmli buffer containing 0.2 mg/ml biotin and analyzed by western blot.

### RNAPII-S2 co-immunoprecipitation

Where indicated, cells were irradiated with UV-C (12 J/m^2^) and harvested 1 h after irradiation. Cells were lysed for 20 min at 4 C in 1 mL EBC-1 buffer (50 mM Tris [pH 7.5], 150 mM NaCl, 2 mM MgCl2, 0.5% NP-40, and protease inhibitor cocktail (Roche)), followed by collection of the chromatin-enriched pellet by centrifugation. Pellets were resuspended in EBC-1 buffer with 500 U/mL Benzonase Nuclease (Novagen) and 2 μg Pol II-S2 (ab5095, Abcam) 1 h at 4 °C. Next, the salt concentration was increased to 300 mM NaCl. Samples were incubated for an additional 30 min at 4 °C, followed by centrifugation at 14.000 x g. 50 μL of supernatant was mixed with 50 μL 2x Laemmli buffer as input sample. The remaining supernatant was incubated with 20 μL equilibrated Protein A agarose beads (Millipore) at 4 °C for 90 min (except for GFP-trap agarose incubated samples). Next, beads were washed 6 times with EBC-2 buffer (50 mM Tris, pH 7.5, 300 mM NaCl, 0.5% NP-40, 1 mM EDTA), and boiled in Laemmli-SDS buffer, followed by analysis by western blot.

**Supplementary Table 1:** Cell lines.

| Cell lines | Source |
| --- | --- |
| RPE1-iCas9 (WT) | 5 |
| RPE1-iCas9 CSB-KO (1-15) | 5 |
| RPE1-iCas9 DOT1L-KO2 | This study |
| RPE1-iCas9 DOT1L-KO6 | This study |
| RPE1-iCas9 KI ELOF1-ultraID-TY #3 | This study |
| RPE1-iCas9 KI ELOF1 T2A-ultraID-TY #4 | This study |
| RPE1-iCas9 KI CSB-KO (1-15) ELOF1-UltraID #2-10 | <sup>5</sup> and this study |

**Supplementary Table 2:**
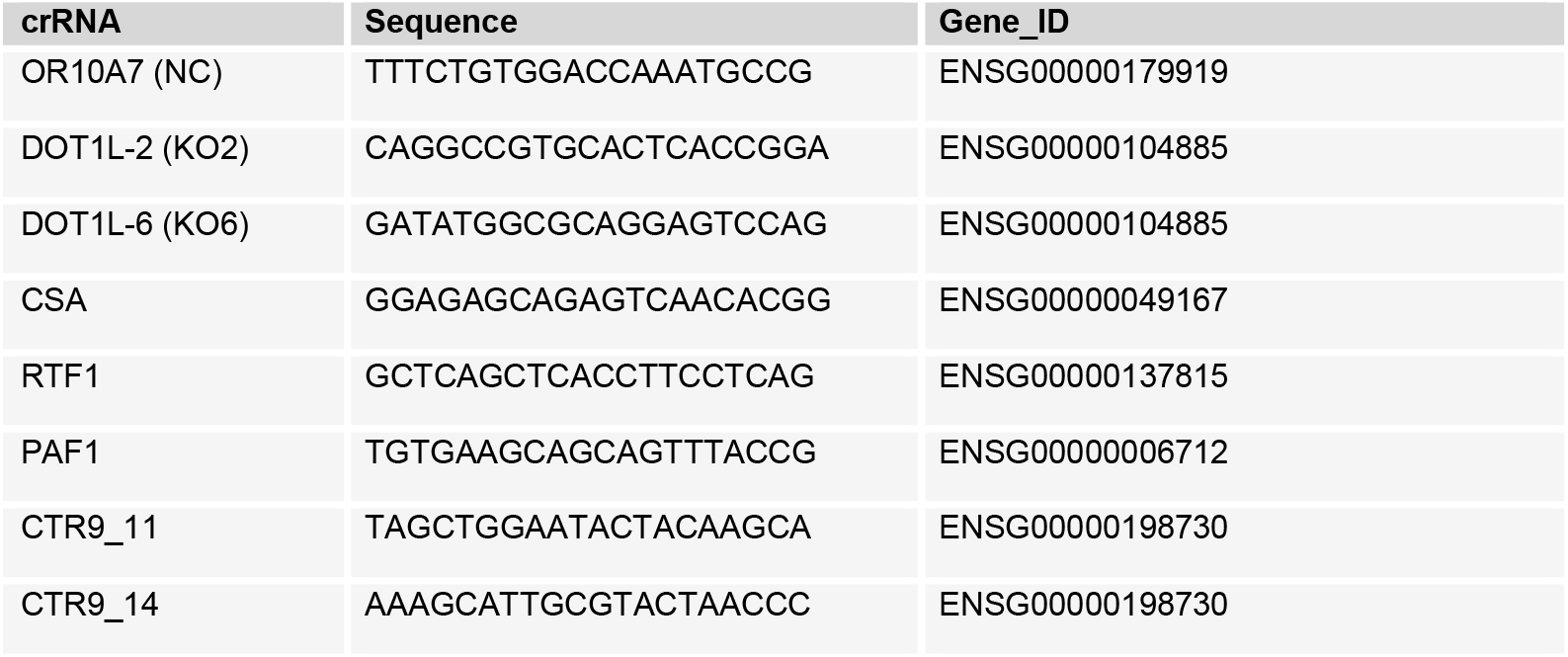
crRNAs.

**Supplementary Table 3:** Plasmids.

| Plasmid | Origin | Identifier |
| --- | --- | --- |
| pML#355_H.Arm1_ELOF1_T2A_ultraID_H.Arm2 | Addgene plasmid #72824 | pML#355 |
| pML#385_H.Arm1_T2A_ELOF_T2A_ultraID_H.Arm2 | Addgene plasmid #72824 | pML#385 |
| pML#187_pX458-(Cas9-2A-GFP)-sgELOF1-3 | Addgene plasmid #48138 | pML#187 |

**Supplementary Table 4:**
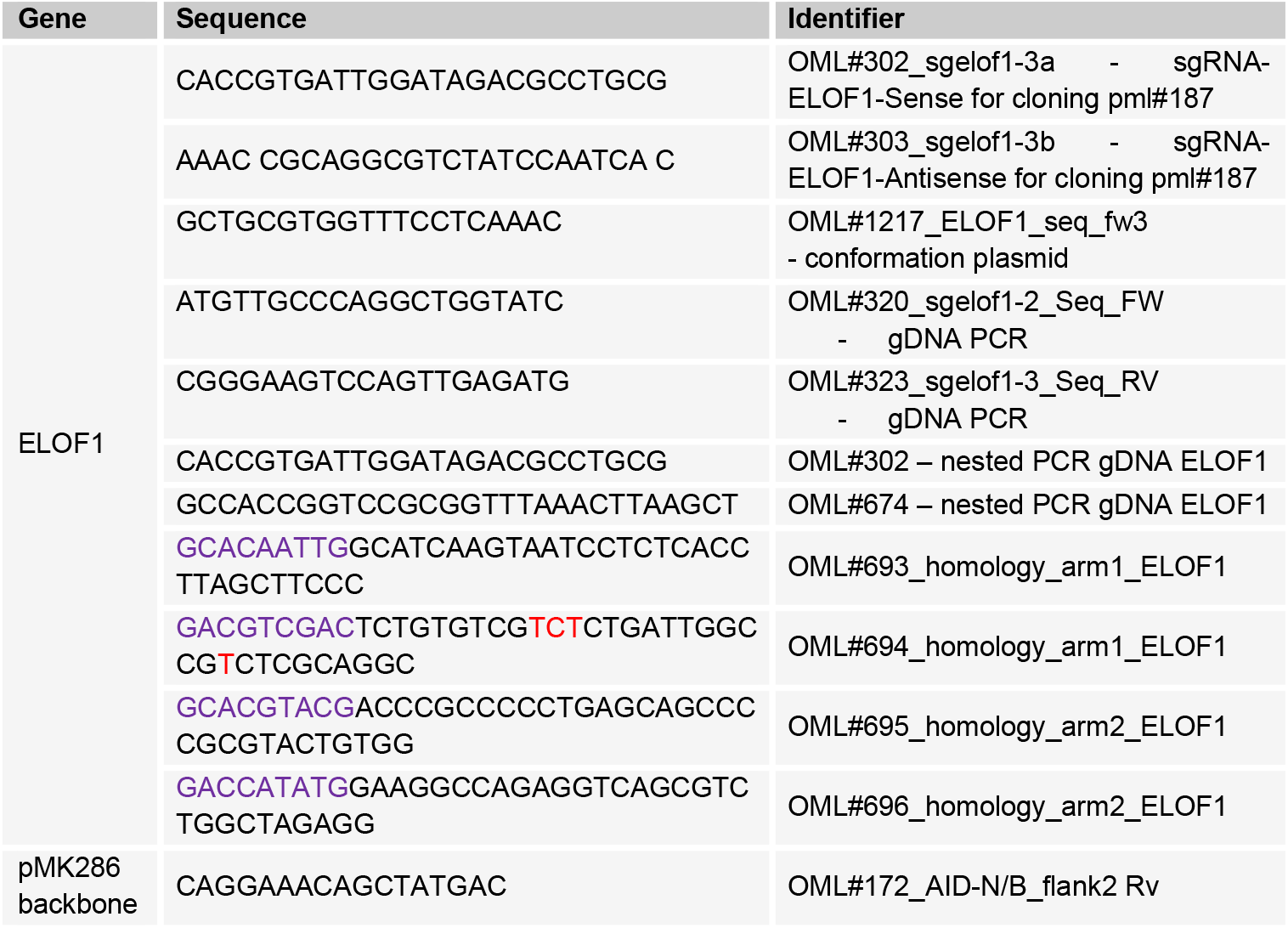
Oligos.

**Supplementary Table 5:** Antibodies.

| Antibody | Host | Company (reference) | Use | Identifier |
| --- | --- | --- | --- | --- |
| CSA/ERCC8 | Rabbit | Abcam, #137033 (EPR9237) | WB: 1:1000 | aML#028 |
| α-Tubulin | Mouse | Sigma, #T6199 (DM1A) | WB: 1:1000 | aML#008 |
| HSPA4 | Rabbit | Novus Biologicals, NBP1-81696 | WB: 1:1000 | aML#114 |
| H3K4me<sub>3</sub> | Rabbit | Cell Signaling, #9751S | WB: 1:1000<br>IF: 1:1000 | aML#237 |
| DOT1L | Rabbit | Cell Signaling, #77087 (D1W4Z) | WB: 1:1000 | aML#216 |
| H3K79me<sub>2</sub> | Rabbit | Abcam, ab3594 | WB: 1:1000 | aML#227 |
| H2B-K120<sub>Ub</sub> | Rabbit | Cell Signaling, 5546S | WB: 1:1000<br>IF: 1:1000 | aML#177 |
| PAF1 | Rabbit | Bethyl, A300-172A | WB: 1:3000 | aML#022 |
| RTF1 | Rabbit | Bethyl, A300-178A | WB: 1:1000 | aML#257 |
| RNAPII | Rabbit | Bethyl, A304-405A | WB: 1:1000 | aML#088 |
| RNAPII-S2 | Rabbit | Abcam, ab5095 | IP: 1:250<br>WB: 1:1000 | aML#024 |
| SPT5 | Mouse | Santa Cruz, sc-133217 | WB: 1:1000 | aML#122 |
| CTR9 | Rabbit | Bethyl, A301-395A | WB: 1:1000 | aML#049 |
| TY1 | Mouse | Diagenode, C15200054 | WB: 1:1000 | aML#154 |
| Rabbit Alexa 647 | Goat | Thermo Fisher, A-21245 | IF: 1:1000 | aML#016 |
| CF770 Goat Anti-Mouse IgG (H+L) | Goat | Biotium, VWR #20077 | WB:<br>1:10000 | aML#009 |
| CF680 Goat Anti-Rabbit IgG (H+L), | Goat | Biotum, VWR#: 20067 | WB:<br>1:10000 | aML#010 |

## Notes

### Competing Interest Statement

The authors have declared no competing interest.

### Summary of Updates

Figures incorporated in manuscript file for easier reading

